# TissueNarrator: Generative Modeling of Spatial Transcriptomics with Large Language Models

**DOI:** 10.1101/2025.11.24.690325

**Authors:** Sizhe Liu, Junjie Tang, Jian Ma, Shaoheng Liang

**Author notes:** Correspondence (S.Liang) and (J.M.).

## Abstract

The intricate spatial organization and molecular communication among cells are fundamental to multicellular systems. Spatial transcriptomics (ST) enables gene expression profiling while preserving spatial context, providing rich data for studying cellular interactions and tissue dynamics. However, most existing computational approaches focus on embedding-based tasks and provide limited generative capacity for simulating cell behavior *in situ*. Moreover, accurately interpreting spatial interactions requires extensive biological knowledge, which current models do not incorporate. Here, we introduce TissueNarrator, a framework that reformulates spatial omics analysis as a language modeling problem. By representing tissue sections as spatial sentences – rank-based gene lists augmented with spatial coordinates and metadata – TissueNarrator leverages pretrained large language models (LLMs) to learn spatially conditioned gene expression patterns. The model generates realistic, context-aware cellular profiles, predicts intercellular interactions, and performs *in silico* perturbation analyses. Across multiple ST technologies (MERFISH, Perturb-FISH, and CosMx SMI), TissueNarrator achieves superior quantitative performance and recovers biologically meaningful ligand–receptor and signaling pathways. Furthermore, a conversational inference mode enables natural-language querying of tissue organization. By integrating pretrained biological knowledge with spatial context, TissueNarrator establishes a new, scalable generative paradigm for modeling, simulating, and reasoning about tissue systems.

## Introduction

In multicellular organisms, cells are arranged in intricate spatial architectures that regulate development, homeostasis, and disease progression [1–3]. The fate and behavior of every cell are tightly modulated by its tissue microenvironment [4]. Understanding intracellular pathways, intercellular signaling, and spatial organization is therefore central to decoding physiological and pathological processes [5]. Spatial transcriptomics (ST) has recently emerged as a transformative technology that measures gene expression while preserving spatial context [6], offering a powerful foundation for studying tissue organization and cell-cell communication [7–9]. However, interpreting ST data requires extensive biological knowledge [10], and current computational approaches remain largely descriptive. Knowledge-based, spatially aware models are needed to infer how gene expression and cell composition change in response to environmental perturbations – a key step toward advancing both cell biology and drug discovery [11–13].

Large language models (LLMs), such as the GPT series [14], have revolutionized artificial intelligence by learning contextual representation through autoregressive prediction on massive text corpora [15]. Recent works like Cell2Sentence (C2S) [16] and Cell-o1 [17] have adapted general-purpose LLMs to biological data, demonstrating that models pretrained on natural language can transfer their biological knowledge to single-cell analysis [18, 19]. LLMs’ ability to integrate large contextual information has also inspired non-language transformer-based models that capture gene-gene interactions in single cells [19–23]. Extensions such as scGPT-spatial [24] and Nicheformer [25] further introduced transformer frameworks for ST data, yet they primarily focus on embedding-based analysis – such as spatial domain identification or cell-type deconvolution [19] – and lack the generative ability to simulate how a cell’s expression profile changes *in situ*. STEAMBOAT [26] further showed that transformer-based architectures can model multiscale cell–cell interactions. Nonetheless, these methods do not process natural language, and thus lack the ability to combine generative modeling with biological priors encoded in pretrained LLMs [18]. As a result, existing frameworks cannot generatively simulate cells *in situ*, thereby limiting realistic predictions of a cell’s behavior in a hypothetically perturbed tissue.

To fill this gap, we introduce TissueNarrator, a knowledge-driven generative framework for simulating spatial biology (**Fig**. 1A). It harnesses the power of LLMs to understand both intra- and intercellular interactions in spatial context. We design “spatial sentences” to represent a piece of tissue by adding spatial coordinates and cell metadata to the textual “cell sentences” of individual cells [16]. In effect, these representations *let cells tell their tissue story* in a form that LLMs can read and extend, without requiring specialized model components. This formulation allows TissueNarrator to unleash the generative capacity of LLMs for predicting new cells in diverse counterfactual tissue environments. In particular, we focus on two generation scenarios across different contexts: inferring the cell type and gene expression of a cell given its neighboring cells and optional metadata. Our evaluation demonstrates that the generated cells capture biologically meaningful patterns, including interactions between non-neuronal cells and immune–cancer cell communication. Beyond cell generation, TissueNarrator supports natural-language querying of tissue structure to facilitate intuitive exploration of spatial organization. Finally, TissueNarrator is designed to be flexible and extensible, supporting diverse tasks through a conversational interface without requiring architectural modifications.

**Figure 1:**
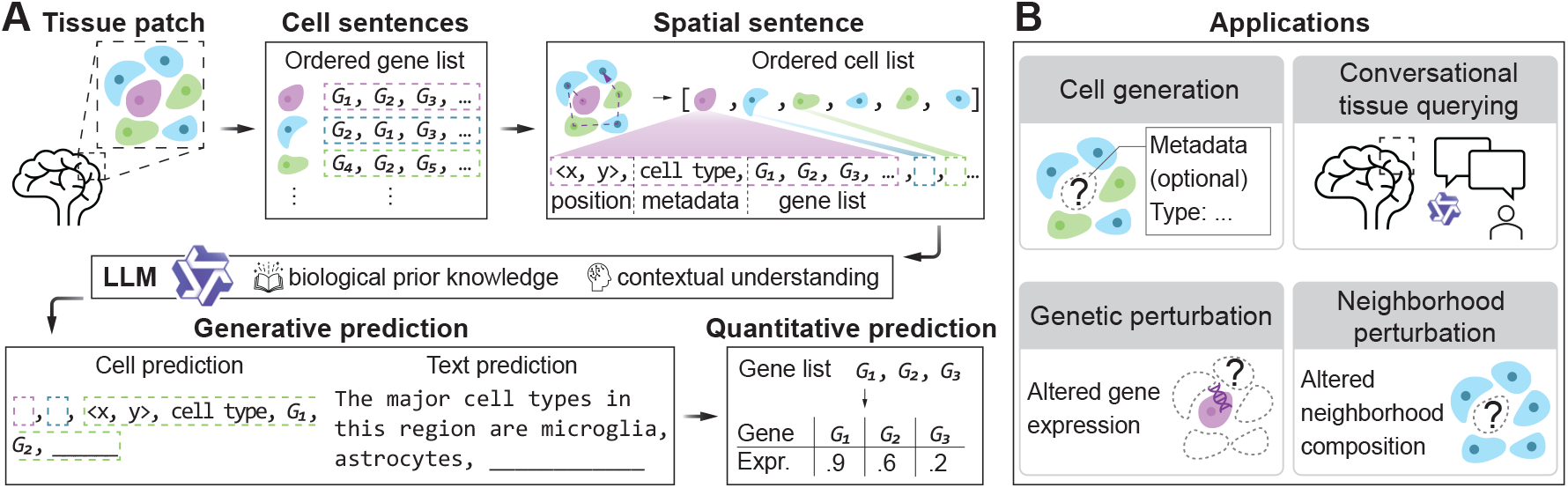
Overview of the TissueNarrator framework. **A.**TissueNarrator takes as input a tissue patch, encodes individual cells into cell sentences, and assembles a spatial sentence from neighboring cells. The spatial sentence can be interpreted by an LLM with biological prior knowledge and contextual understanding to predict gene expressions in a cell and to answer questions about tissues. Text output of the LLM can be further transformed into gene expression levels. **B**. TissueNarrator supports four downstream tasks: cell generation, neighborhood perturbation, genetic perturbation, and conversational tissue querying. For cell generation tasks, model-generated cell sentences can be converted back into gene expression space.

## Results

### Overview of generative modeling in TissueNarrator

TissueNarrator reformulates spatial transcriptomics modeling as a conditional generative problem, bridging the gap between biological systems and linguistic reasoning. This requires resolving two fundamental incompatibilities: the dimensionality mismatch of mapping 2D tissues to 1D sequences, and the semantic gap in spatial representation. We address the former via a topology-preserving serialization strategy that maintains local neighborhood context. For the latter, we demonstrate that explicit numerical tokens offer a superior alternative to learnable positional embeddings, bypassing the need for de novo representation learning by activating the LLM’s pretrained geometric priors. These innovations converge in what we call a “spatial sentence” – an expressive, scalable representation of a tissue that encodes complex cellular environments as structured natural language. This format enables effective fine-tuning of an LLM on spatial transcriptomics data without specialized model architectures.

An overview of the TissueNarrator framework is shown in **Fig**. 1A. On an input tissue section, TissueNarrator starts by transforming each cell into the natural language space by encoding gene expression as “cell sentences”, a list of genes ranked by gene expression level. All cell sentences in a tissue patch are then assembled into a 1D “spatial sentence”, where cells are serialized by proximity-based traversal. This spatial sentence then serves as the input for fine-tuning an LLM to learn cell generation through next-token prediction. The fine-tuned model fits multiple downstream applications, including cell generation, neighborhood perturbation, genetic perturbation, and conversational tissue querying. Further details of TissueNarrator can be found in the **Methods** section.

We evaluate our model on three spatial transcriptomics datasets that cover different tissue types (mouse brain, human melanoma, and ovarian cancer) sequenced by different ST technologies (MER-FISH, Perturb-FISH, and CosMx SMI):

- **Whole Mouse Brain (MERFISH)** [27] includes 129 adult mouse brain sections with 2.6 million cells profiled by MERFISH using a 1,122-gene panel. Cells are annotated into 35 major types and mapped to anatomical regions provided by the Allen Institute. *Tasks:* Cell generation, Neighborhood perturbation, Conversational tissue querying.
- **Human melanoma (Perturb-FISH)** [13] includes 35 gene perturbations performed in A375 human melanoma cells. Expression of 500 genes from an immune-oncology panel was measured to capture transcriptional responses to these perturbations within the tumor microenvironment. *Task:* Genetic perturbation.
- **Human ovarian cancer (CosMx SMI)** [28] includes high-grade serous tubo-ovarian cancer samples. We focus on the untreated subset consisting of 27 adnexa and 21 omentum samples, each with 2,000-10,000 cells and 979 genes measured. *Task:* Cell generation (immune–cancer interaction).

### TissueNarrator generates cells that capture realistic cell-cell interaction patterns

The gene expression of every cell is modulated by its tissue microenvironment. Modeling these context effects is therefore an important test of our generative approach. Here, we demonstrate TissueNarrator’s core ability to model these interactions through context-aware cell generation. We first used the MERFISH mouse brain dataset [27], which provides detailed annotations of cell types across multiple brain regions (**Fig**. 2A). Sections were randomly divided into training, validation, and test sets in an 8 : 1 : 1 ratio.

**Figure 2:**
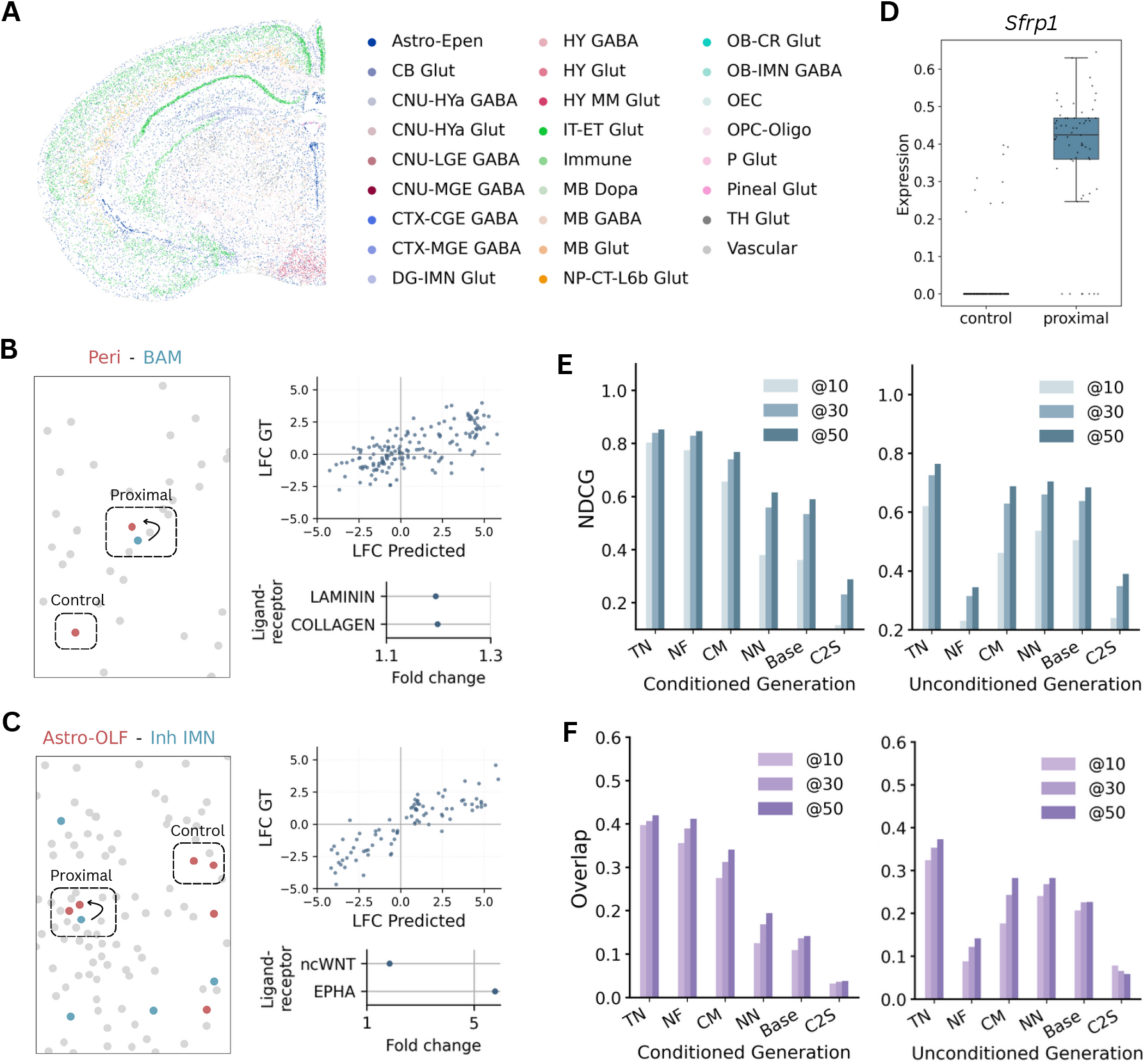
Cell-cell interaction modeling in mouse brain MERFISH data. **A**. *In situ* plot of annotated cell types in a representative slide. **B**. Left: pericytes (Peri; red) are designated as proximal and control depending on their distance to the closest border-associated macrophage (BAM; blue). Top-right: correlation of log-fold changes between generated proximal vs control and ground-truth proximal vs control comparisons. Bottom- right: ligand—receptor pathways upregulated in proximal vs control cell pairs. **C**. Same as B, but for olfactory astrocytes (Astro-OLF; red) and inhibitory immature neurons (Inh IMN; blue). **D**. Generated expression of *Sfrp1*, a gene previously reported to be upregulated in Astro-OLF cells proximal to Inh IMN cells. **E–F**. NDCG (E) and Overlap Score (F) comparing model-generated and ground-truth cell sentences under two evaluation settings. The left panels show the conditioned generation results, and the right panels show the unconditioned generation results. “@k” indicates evaluation using the top 10, 30, or 50 genes. TN, TissueNarrator; NF, Nicheformer; CM, Class Mean; NN, Nearest Neighbor; C2S, C2S-Scale.

Brain function relies on neuronal circuits and the intricate support networks of non-neuronal cells [27,29]. Investigating these interactions allows us to validate TissueNarrator across different biological contexts. Hence, we performed two case studies to test whether TissueNarrator could recover known intercellular interactions [27] from held-out test sections, focusing on interactions crucial for brain functions.

For non-neuron interactions, we investigated the interaction between pericytes and border-associated macrophages (BAMs). Pericytes within 30 *µ*m of BAMs were labeled as proximal, and distant pericytes were labeled as controls. We then generated pericytes conditioned on their neighbors (**Methods**), masking all neighboring pericytes to avoid direct information leakage. Differential expression analyses between the generated proximal and control cells showed strong agreement with ground-truth log-fold changes (Spearman *ρ* = 0.67, **Fig**. 2B). Ligand–receptor analysis using CellChat [30] found significant upregulation of collagen and laminin pathways in model-generated proximal pericytes. This was partially consistent with the ground truth, which had upregulated laminin (the model additionally highlighted a collagen involvement).

For neuron–non-neuron interactions, we examined astrocytes and inhibitory immature neurons (IMNs). TissueNarrator accurately generated astrocytes depending on the presence of IMNs and agreed with ground-truth log-fold changes (Spearman *ρ* = 0.87), recovering known ncWNT and EPHA signaling pathways (**Fig**. 2C). Notably, the original study [27] reported that *Sfrp1*, a WNT signaling modulator, was upregulated in astrocytes near IMNs. TissueNarrator successfully recovered this differential expression pattern (**Fig**. 2D), whereas none of the baseline methods did.

Finally, to quantitatively evaluate TissueNarrator’s cell generation ability across all cell types, we predicted the gene expression profile of a query cell using its cell type and the gene expression of its neighboring cells as input. For a more stringent evaluation, same-type neighbors were intentionally excluded (cross-type generation). This setup, applied to sampled cells from all classes (up to 500 pertype), provided a robust test of intercellular communication modeling. TissueNarrator achieved the highest NDCG and overlap scores (**Methods**) across all baselines (**Fig**. 2E-F, left panels), indicating the most accurate ranking and identification of biologically relevant genes. Furthermore, TissueNarrator maintained high accuracy even when the query cell’s type was not provided (while same-type neighbors were included) (**Fig**. 2E–F, right). An ablation analysis (**Supplementary Note** F) further confirmed that spatial coordinate information, metadata, neighbor context, and the traversal strategy each contribute substantially to model performance.

In summary, these results demonstrate that TissueNarrator effectively learns the molecular grammar of tissue organization, allowing it to reconstruct cell states and decipher intercellular communication patterns from spatial transcriptomics data.

### TissueNarrator enables efficient *in silico* neighborhood perturbation modeling

Perturbation studies are crucial for elucidating how cells adapt to changes in their environment or genotype, offering insight into disease mechanisms and potential therapies [11]. In spatial contexts, two representative forms of perturbation are particularly informative: (1) altering the cell-type composition of a cell’s neighborhood (e.g., adding or removing a specific cell type), and (2) introducing a genetic modification in a cell, such as gene knockouts in individual cells, and observing their propagated effects on neighboring cells. However, such experiments *in vivo* are time-consuming and costly. Tissue-Narrator addresses this challenge by enabling *in silico* spatial perturbation modeling, allowing rapid generation of novel and testable hypotheses.

We first demonstrated TissueNarrator’s ability to model the effects of altering neighborhood cell-type composition using the MERFISH mouse brain dataset. Olfactory ensheathing cells (OEC) and thalamus glutamatergic neurons (TH Glut) are two cell types that normally reside in distinct brain regions and are not typically found in proximity. As a counterfactual experiment, we performed an *in silico* transplant by placing TH Glut cells near OECs and examined the resulting changes in OEC gene expression (**Fig**. 3A). Gene Set Enrichment Analysis (GSEA) on the generated OECs revealed significant downregulation of extracellular matrix (ECM) pathways (**Fig**. 3B). This finding is consistent with previous observations that OECs participate in ECM remodeling [31] and supports the biologically plausible hypothesis that the presence of transplanted TH Glut cells could trigger ECM remodeling in nearby OECs, reflecting their adaptive response to altered neuronal signaling.

**Figure 3:**
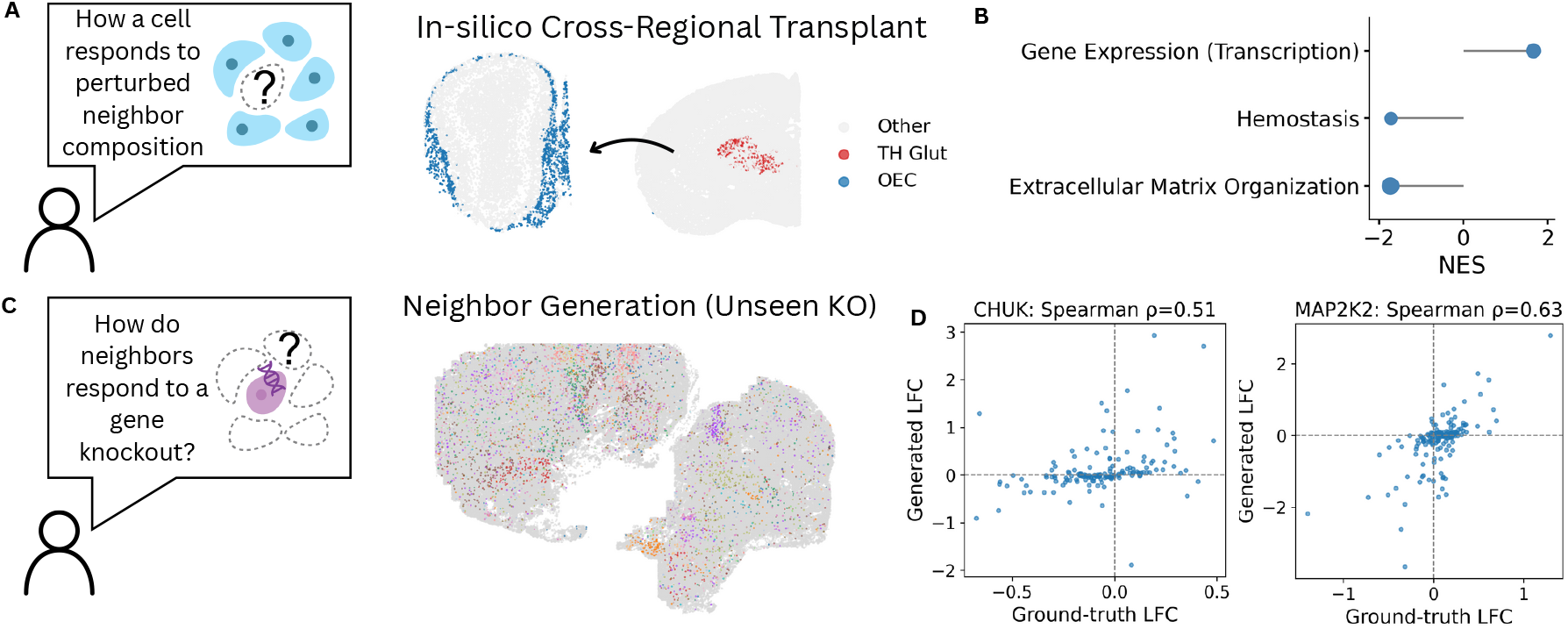
Modeling spatial and genetic perturbations *in silico*. **A**. Scatter plot of the *in silico* cross-regional transplant experiment. A thalamus glutamatergic neuron (TH Glut; red) from one section is transplanted into another section, near olfactory ensheathing cells (OEC; blue) **B**. GSEA results for OEC cells proximal to the transplanted TH Glut cell, showing downregulation of ECM pathways. **C**. Visualization of melanoma tumor cells from the Perturb-FISH dataset, including 35 gene perturbations. Each perturbation is shown in a unique color, while grey cells represent unassigned cases. **D**. Comparison of log-fold changes between generated and ground-truth differential expression for proximal knockout vs control neighbors. Knock-out of *CHUK* and *MAP2K2* are shown as examples.

We next assessed TissueNarrator’s capability to model the propagation from genetic perturbations in one cell to its neighbors. For this, we used the Perturb-FISH dataset of melanoma tumor cells [13], encompassing 35 gene perturbations (**Fig**. 3C). Given the limited number of perturbations, we adopted a leave-one-out cross-split strategy: when testing one perturbation, its cells were held out, with the remainder used for training. During inference, TissueNarrator was given a single unseen perturbed cancer cell and tasked with generating the gene expression of its neighboring T cells. We then performed differential expression analysis, comparing generated T cells near control cells to those near unseen knockout cells. The log-fold changes between generated and ground-truth data showed strong agreement (**Fig**. 3D). For example, TissueNarrator accurately captured the magnitude of perturbation effects, producing smaller LFCs for *CHUK* knockouts and larger ones for *MAP2K2* (both with Spearman *ρ >* 0.5).

These results demonstrate TissueNarrator’s unique ability to predict neighborhood-level perturbation effects from a single perturbed cell input.

### TissueNarrator models immune infiltration programs in spatial contexts

Tubo-ovarian high-grade serous carcinoma (HGSC) is the most common and aggressive form of ovarian cancer. Understanding the interaction of immune and malignant cells in the tumor microenvironment (TME) is essential for elucidating disease progression and therapeutic response (**Fig**. 4A) [28, 32, 33]. To showcase TissueNarrator’s ability to capture these spatial immune–tumor relationships, we focus on the tumor infiltration program (TIP), a set of genes that are significantly overexpressed or underexpressed in specific immune cell subtypes depending on their proximity to malignant cells (**Fig**. 4B).

Using the cross-type generation setting, with immune cells as prediction targets, we calculated AU-ROC scores on known immune subtype gene signatures to evaluate how well predicted cells matched their expected subtypes *in situ*. We observed AUROC scores greater than 0.67 in the most abundant immune subtypes (monocytes, B cells, and CD8^+^ T cells; **Fig**. 4C). While performance was more modest in rarer subtypes, all immune subtypes consistently showed AUROC scores greater than 0.5. These results underline TissueNarrator’s ability to disentangle genes specific to immune subtypes within the TME, confirming that it captured distinct immune identities rather than simply reverting to an average gene expression profile.

Next, to probe the TIP captured by TissueNarrator, we performed differential expression analysis for each immune subtype, comparing predicted immune cells located near tumor regions with those farther away. Many of the genes identified by TissueNarrator aligned with those reported in [28]. For example, in CD8^+^ T cells, TissueNarrator accurately reported upregulation of *CCL3, CTSW, HSP90B1*, etc., alongside downregulation of *CXCR4, CD28, CD44*, etc. GSEA further highlighted significant enrichment of the CD28-related pathway, indicating that the predicted results captured specific T-cell signaling (**Fig**. 4D). Similarly, in monocytes, TissueNarrator accurately identified upregulation of *CLEC5A, SPP1, SRGN*, etc., and downregulation of *CLEC10A, IL2RA, LYZ*, etc. (**Fig**. 4E).

Together, these results show that TissueNarrator not only models immune cell identity, but also accurately captures their context-dependent functional states in the TME.

**Figure 4:**
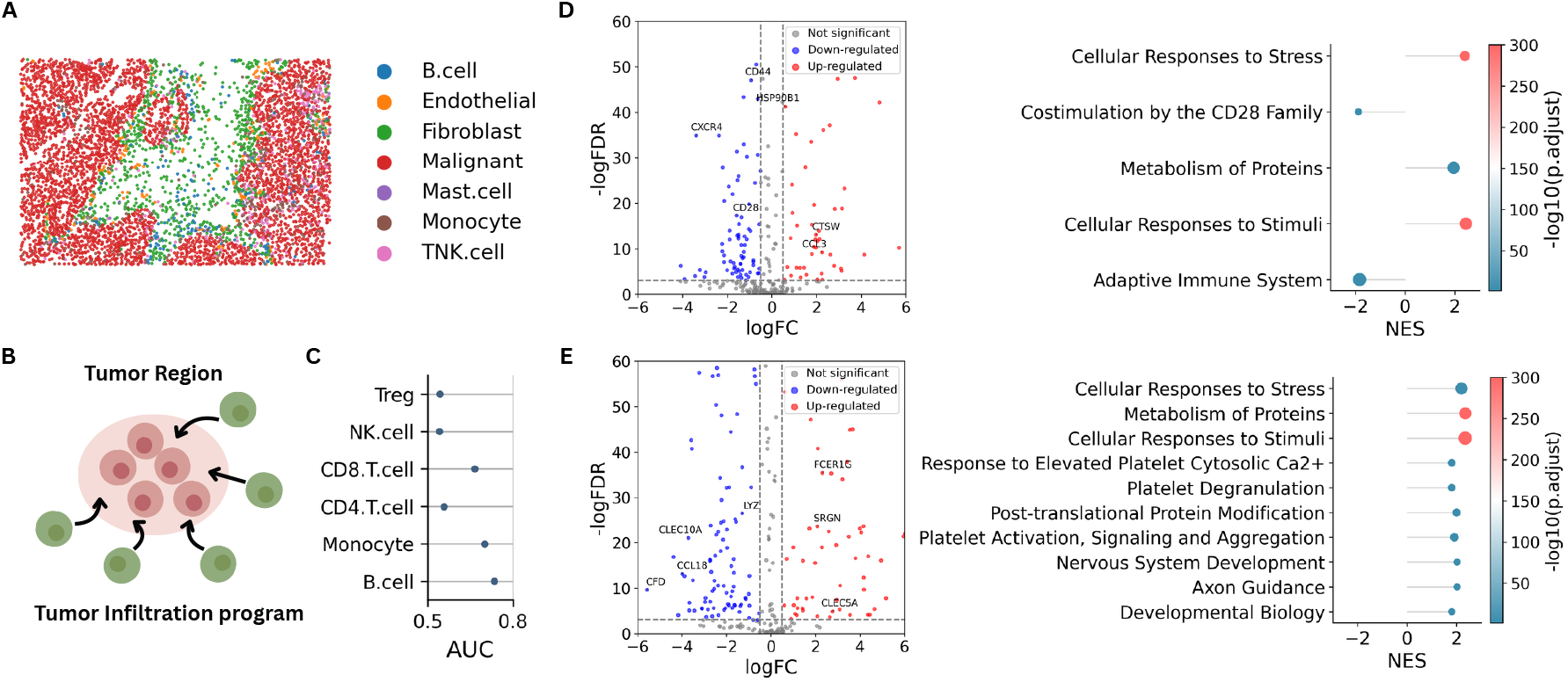
Modeling immune–tumor interactions in ovarian cancer. **A**. Spatial distribution of cell types in a representative sample (SMI_T10_F001). **B**. Illustration of the tumor infiltration program, showing genes that are differentially expressed in immune cells as a function of their proximity to malignant cells within the tumor region. **C**. Area under the ROC curve (AUC) scores computed from the expression of reported marker genes for each immune subtype. **D**. Differential expression of model-generated CD8+ T cells proximal vs non-proximal to malignant cells. Significantly differentially expressed genes that align with those reported in the original study are highlighted in the volcano plot on the left (thresholds: |logFC| ≥ 0.5, FDR ≤ 0.001), and GSEA results are shown on the right. **E**. Same analysis as in D, but for monocytes.

### TissueNarrator enables interactive Q&A about spatial regions

To evaluate TissueNarrator’s ability to apply biological prior knowledge and learned spatial knowledge in a Q&A setting, we constructed a proof-of-concept dataset covering three tasks: given the name of an anatomical structure, (1) return the top 5 most abundant cell types, (2) list the top 20 highly expressed genes, and (3) generate a natural-language description of the structure. Details of data curation and task examples are provided in **Supplementary Note** D. We split the dataset by structure name, fine-tuned TissueNarrator, and evaluated the model on questions derived from held-out structures. TissueNarrator generalizes to unseen anatomical structure names in natural language queries by building on the pretrained knowledge of its LLM backbone. For example, it can generate the exact top five cell types for the Dorsal peduncular area (**Fig**. 5A) and also produces a coherent, biologically grounded paragraph describing cranial nerves (**Fig**. 5B), These examples highlight the model’s ability to link spatial context with natural language explanations, a bridge between data-driven cell modeling and user-friendly querying. Moreover, fine-tuning TissueNarrator outperformed directly fine-tuning `Qwen-4B-Base`. This suggests that the spatial sentence training stage not only teaches the model cell generation, but also enables a deeper spatial understanding that transfers to Q&A tasks (**Supplementary Note** D).

**Figure 5:**
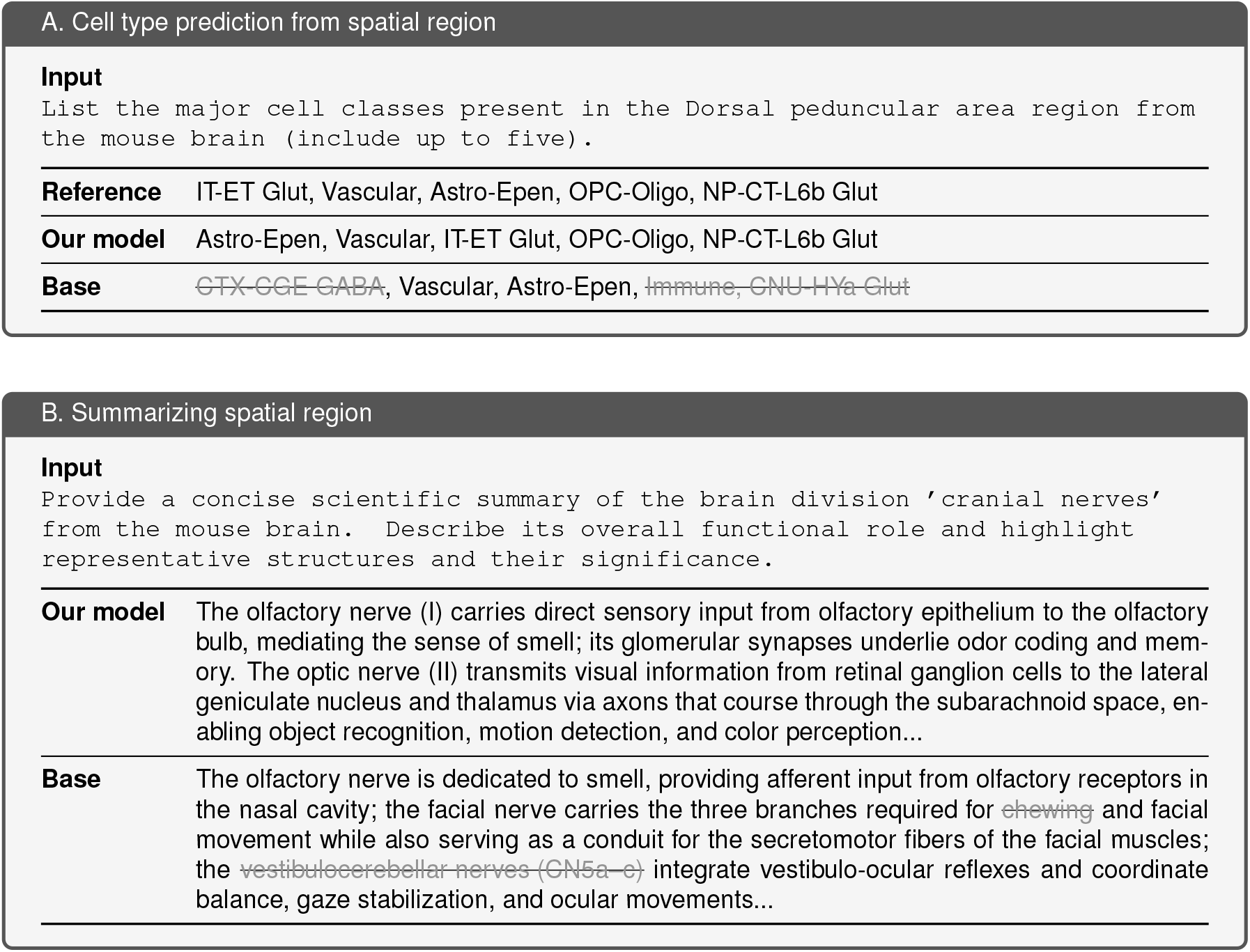
Case studies of TissueNarrator in spatial Q&A tasks. **A**. Cell type prediction in the Dorsal peduncular area. Incorrect content is marked using gray strike-through. **B**. Region summarization for cranial nerve structures.

## Discussion

In this work, we introduced TissueNarrator, an expressive, scalable framework that effectively adapts LLMs to modeling spatial transcriptomics by integrating biomedical prior knowledge with spatial context. By converting spatial measurements into textual sequences, TissueNarrator leverages the extensive knowledge and contextual reasoning capabilities of LLMs to provide deeper insight into both intra- and inter-cellular interactions. In essence, we recast a spatial biology problem into a language problem, allowing us to harness an LLM’s power in a novel way. Notably, we demonstrate that LLMs can effectively interpret (1) explicit numerical spatial coordinates and (2) cell sequences ordered via topology-preserving serialization. This finding establishes a streamlined, text-native alternative to more computationally involved graph-based encoding of spatial data.

A central conceptual contribution of TissueNarrator is reframing ST analysis as a language modeling problem. Existing spatial models focus primarily on static or embedding-based representations and lack generative capability for producing context-aware cellular profiles. In contrast, TissueNarrator learns to simulate realistic cellular profiles conditioned on their spatial environment, enabling exploration of unseen spatial arrangements and *in silico* perturbations. This framing shifts ST analysis from a descriptive or predictive paradigm to a generative one, where models can simulate plausible tissue states under hypothetical scenarios. Such virtual experimentation can complement biological studies by narrowing hypothesis spaces and prioritizing conditions for validation.

TissueNarrator also highlights several directions for future improvement and expansion. Its expressive capacity and ability to generalize across diverse datasets are currently constrained by model size and the scope of training data. Scaling model capacity and increasing dataset diversity would allow learning across diverse tissue types and spatial technologies. Larger models may also unlock more advanced capabilities, such as chain-of-thought reasoning, which could help provide mechanistic interpretations of generated results and further integrate biological prior knowledge into complex tasks. In addition, current LLM context length limits TissueNarrator to neighborhoods containing only hundreds of cells, making it challenging to represent tissue phenomena involving many cells or long-range spatial dependencies. Architectural advances or improved context-selection strategies could extend effective context length and enable modeling of multi-scale tissue organization.

In conclusion, TissueNarrator represents an important step toward developing scalable models capable of simulating cellular organization and responses within complex tissue systems. We envision TissueNarrator and related approaches evolving into general-purpose frameworks for spatial biology and helping researchers “narrate” tissues, guiding experimental design, generating testable hypotheses, and accelerating discovery across tissue architecture, disease mechanisms, and therapeutic development.

## Methods

The TissueNarrator framework first encodes individual cells as “cell sentences” and then assembles neighboring cells into “spatial sentences”. The natural-language spatial sentences are then used to fine-tune an LLM to learn cell and text generation through next-token prediction. TissueNarrator enables multiple downstream applications, including cell generation, neighborhood perturbation, genetic perturbation, and conversational tissue querying. The details are as follows.

### Cell sentence transformation

We extended the rank-based gene representation introduced in Cell2Sentence (C2S) [16] to provide a compact and expressive format for LLM-based analysis of spatial data. While positional encoding is an established approach for incorporating cell locations [25, 34], studies in other fields of machine learning [35, 36] and our empirical observations suggest that its effectiveness can be limited when the model is not extensively pretrained on spatial signals. Instead, we directly use spatial coordinates (*x*_*i*_, *y*_*i*_) as numerical tokens within the sentence. This design aligns with emerging evidence that LLMs can inherently process and make sense of numerical coordinates [37], providing a more direct and interpretable representation of spatial information. Specifically, in TissueNarrator each cell is represented as a cell sentence (**Fig**. 1A):

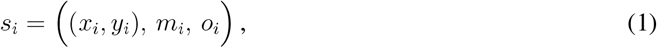

where (*x*_*i*_, *y*_*i*_) denotes its spatial coordinates, *m*_*i*_ represents optional metadata (e.g., cell type or spatial domain), and *o*_*i*_ is the ranked gene sentence, defined as a list of gene symbols ordered by decreasing transcript abundance [16].

**Spatial sentence transformation**

For each tissue section, we segmented the entire section into square regions of size *L* × *L* micrometers, referred to as spatial neighborhoods. For each spatial neighborhood *j*, cells with their cell sentences 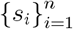 were assembled in pre-determined spatial order to construct one spatial sentence (**Fig**. 1A) representing the local cellular composition and organization:

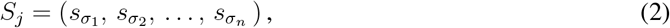

where *σ* = (*σ*_1_, *σ*_2_, …, *σ*_*n*_) denotes the traversal order of cells within the frame.

Although ST data capture cells in a spatial tissue context, they lack an inherent sequential structure, whereas language models require ordered token inputs. To bridge this gap, we introduce an artificial traversal order that serializes the spatial arrangement of cells into a sequence suitable for LLM processing. This design is partially inspired by GeST [38], with two key improvements. First, the traversal weights neighboring cells more heavily, preserving local spatial continuity so that each preceding cell provides informative spatial and transcriptional context for the next. Second, controlled randomness improves robustness and prevents overfitting to specific spatial patterns.

Traversal begins by randomly selecting one of five anchor points – the geometric center or the four corners. Let *a* denote the chosen anchor. Cells are then sampled sequentially from the unvisited set 𝒰 with distance-weighted probability:

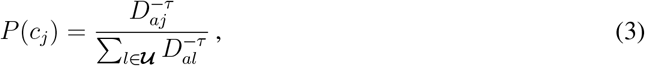

where *D*_*aj*_ is the Euclidean distance between cell *j* and the anchor, and *τ* (default *τ* = 1) controls randomness. This process continues until all cells in the neighborhood have been visited, yielding the traversal sequence *σ*. Examples of spatial sentences are provided in **Supplementary Note** A.

### TissueNarrator training

#### Training objective

TissueNarrator is fine-tuned from a pretrained LLM using the standard next-token prediction objective [39], which maximizes the conditional probability of the text sequence.

Each spatial sentence *S*_*j*_ is tokenized into tokens *w*_1_, *w*_2_, …, *w*_*T*_, where *w*_*t*_ denotes the token at position *t*. The language modeling objective is expressed using the cross-entropy loss:

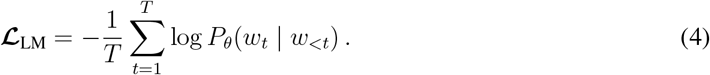

Conceptually, at the intracellular level, the model learns to predict the next gene token within a ranked gene list, while at the intercellular level, it learns to predict the next cell conditioned on the representations of preceding cells.

#### Training protocol

We fine-tune the `Qwen3-4B-Base` model [40] using the Unsloth framework [41]. The model is trained for 3 epochs with a batch size of 2 and a gradient accumulation factor of 4. We use a linear warm-up over the first 1% of steps. To improve efficiency, we adopt Low-Rank Adaptation (LoRA) [42] for parameter-efficient fine-tuning. A context length of 32k tokens is used to accommodate long spatial sentences. Detailed training hyperparameters are provided in **Supplementary Note** B.

### TissueNarratorinference

#### Cell generation mode

Given the cell sentences from neighboring cells and the spatial location of a target cell, TissueNarrator can predict the target cell sentence in two distinct modes, each serving distinct biological purposes. (1) In the conditional generation mode, users can specify the target cell type or other metadata (e.g., tissue region or disease state), enabling controlled exploration of how a particular cell type behaves under a defined spatial context. (2) In the unconditional generation mode, TissueNarrator infers the target cell’s identity and expression profile solely from neighboring spatial cues, reproducing natural spatial distributions without prior knowledge of the target cell type.

#### Quantitative gene expression prediction

To reconstruct quantitative expression values of single cells from the predicted ranked gene sentence output by TissueNarrator, we fit a linear model for each dataset *d*, describing the relationship between a gene’s expression value *e*_*i*_ and its within-cell rank *r*_*i*_:

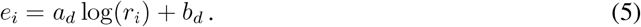

The fitted parameters *a*_*d*_ and *b*_*d*_ are then used to convert predicted gene ranks back into quantitative expression estimates. We demonstrate the robustness of this transformation in **Supplementary Note** C. Through this formulation, TissueNarrator can output results in both ranked-list and quantitative expression formats, supporting analysis in either representation.

#### Extended task capabilities

Following large-scale training on spatial sentences, TissueNarrator can be fine-tuned on smaller, task-specific datasets to extend its functionality beyond cell generation. As a proof of concept, we evaluated this capability using an interactive Q&A task (**Supplementary Note** D), where the model answers descriptive questions about spatial omics. This demonstrates that TissueNarrator can use its pretrained biological knowledge to support interpretable and accessible exploration of tissue structure.

### Real data applications

TissueNarrator generates cells by leveraging a wide range of information. While the location of the cell and the information from neighboring cells are essential, TissueNarrator can also condition on optional metadata such as cell type. This makes TissueNarrator a flexible framework for multiple downstream applications. Specifically, we evaluate TissueNarrator on four key applications (**Fig**. 1B):

- **Missing cell generation:** TissueNarrator predicts a target cell missing from ST using its neighborhood. In particular, it handles the challenging case in which no similar cell types are present nearby.
- **Predicting response to neighborhood composition:** TissueNarrator predicts a target cell within a neighborhood with counterfactual cell-type composition, enabling *in silico* perturbations such as adding or removing neighbor cells.
- **Predicting effects of genetic perturbation:** TissueNarrator predicts a target cell given an edited or perturbed neighboring cell, expanding known single-cell perturbation effects to tissue-level responses.
- **Conversational tissue querying:** TissueNarrator answers spatial omics questions in natural language, enabling intuitive exploration of spatial organization.

### Performance evaluation

#### Quantitative evaluation

As quantitative evaluation metrics, we use the Normalized Discounted Cumulative Gain (NDCG) [43] and the Overlap Score. NDCG evaluates ranking quality by comparing the predicted and ideal gene orders, while the Overlap Score captures how well the model recovers the most relevant genes. For a ranked list of genes, let rel_*i*_ denote the relevance score at rank *i* (set to 1 if the gene is in the ground truth set and 0 otherwise). The Discounted Cumulative Gain (DCG) and Normalized DCG (NDCG) at cutoff *k* are defined as:

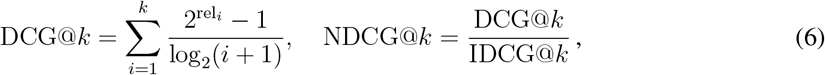

where IDCG@*k* denotes the DCG of the ideal ranking sorted by true relevance.

The Overlap Score computes agreement between the top-*k* predicted and ground-truth genes:

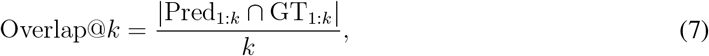

where Pred_1:*k*_ and GT_1:*k*_ denote the sets of the top-*k* predicted and ground-truth genes, respectively. We report results at *k* = 10, 30, and 50, corresponding to NDCG@*k* and Overlap@*k* using the top *k* genes.

#### Qualitative evaluation

Using the reconstructed expression profiles, we perform qualitative evaluations including differential expression (DE) analysis and ligand–receptor interaction analysis to assess the biological plausibility of generated cells.

#### Baseline methods

Nearest Neighbor selects the cell in the prompt that has the smallest Euclidean distance to the target cell. Class Mean reports the mean expression profile of all cells belonging to the same class as the target cell in the training set. Nicheformer is a transformer-based model pretrained on SpatialCorpus-110M, a curated collection comprising over 57 million dissociated and 53 million spatially resolved cells across 73 tissues [25]. We compute cell embeddings and use an MLP layer to predict target gene expression. C2S-Scale is the latest model in the Cell2Sentence series, trained on over 57 million single cells [44]; we use `C2S-Scale-2B` for zero-shot inference. Base refers to the pretrained `Qwen-4B-Base` model [40] from which TissueNarrator is fine-tuned.

For a fair comparison with TissueNarrator in the conditioned generation setting, each baseline is further modified to incorporate metadata, as described in **Supplementary Note** E.

## Code availability

The source code implementing the TissueNarrator framework is available at: https://github.com/Steven51516/tissuenarrator.

## Acknowledgment

This work was supported, in part, by National Institutes of Health Common Fund 4D Nucleome Program grant UM1HG011593 (J.M.); National Institutes of Health Common Fund Cellular Senescence Network Program grant UH3CA268202 (J.M.); and National Institutes of Health grants R01HG007352 (J.M.), R01HG012303 (J.M.), R21DA061481 (J.M.), and R03OD039980 (J.M.). J.M. was additionally supported by the Ray and Stephanie Lane Professorship, a Guggenheim Fellowship from the John Simon Guggenheim Memorial Foundation, and a Google Research Award. S.Liang is a Lane Fellow. The funders had no role in study design, data collection and analysis, decision to publish or preparation of the manuscript.

## Author Contributions

Conceptualization, S.Liu, J.M., S.Liang; Methodology, S.Liu, J.M., S.Liang; Software, S.Liu, S.Liang; Investigation, S.Liu, J.T., J.M., S.Liang; Writing, S.Liu, J.M., S.Liang; Funding Acquisition, J.M.

