## Supplementary Material for "TissueNarrator: Generative Modeling of Spatial Transcriptomics with Large Language Models"

### (SUPPLEMENTARY NOTE)

#### A Spatial Sentence Examples

A. Spatial sentence (without metadata)

**Example**

```
<pos> X: 3798, Y: 3813 <cs> IGFBP6 GJA1 SLC7A11 ITIH5 FN1 IGF2 ...  CPNE8
</cs> <pos> X: 3782, Y: 3852 <cs> GFAP GJA1 NBL1 FNDC1 ACSBG1 ...  CBLN4
</cs> ...
```

B. Spatial sentence (with metadata)

**Example**

```
<pos> X: 3798, Y: 3813 <meta> class:  Vascular <cs> IGFBP6 GJA1 SLC7A11 ITIH5
FN1 IGF2 ...  CPNE8 </cs> <pos> X: 3782, Y: 3852 <meta> class:  OEC <cs> GFAP
GJA1 NBL1 FNDC1 ACSBG1 ...  CBLN4 </cs> ...
```

**Figure S1:** Examples of spatial sentences used for text-based representation of spatial transcriptomics data. **A.** A spatial sentence constructed only from gene expression and spatial coordinates. **B.** A spatial sentence enriched with metadata such as cell class.

#### B Model Hyperparameters

**Table S1:** Training and LoRA hyperparameter settings used in this study.

| **Parameter** | **Setting** |
|---|---|
| *Training* | |
| Max sequence length | 32,000 |
| Training epochs | 3 |
| Batch size per device | 2 |
| Gradient accumulation steps | 4 |
| Learning rate | 2e–4 |
| Warmup ratio | 0.01 |
| Weight decay | 0.01 |
| LR scheduler | `cosine` |
| *LoRA configuration* | |
| Rank ($r$) | 32 |
| LoRA alpha | 32 |
| LoRA dropout | 0 |
| Bias | `none` |
| Target modules | `q_proj, k_proj, v_proj, o_proj, gate_proj, up_proj, down_proj` |
| Gradient checkpointing | `unsloth` |

#### C Robustness of Expression Reconstruction

Although Cell2Sentence [1] demonstrated that quantitative expression values can be robustly reconstructed from ranked representations in single-cell RNA-seq data, spatial transcriptomics data exhibit distinct characteristics beyond simply adding a spatial dimension to single-cell measurements [2]. We therefore further evaluate the robustness of this rank-to-expression transformation in the spatial context.

We benchmarked reconstruction performance using the MERFISH Mouse Brain dataset [3], assessing the ability to recover quantitative gene expression values from ranked gene representations. The reconstructed expression achieved an $R^2$ of 0.91 and a Spearman correlation of 0.95 with the original values, indicating highly consistent recovery. In addition, the UMAP visualization qualitatively shows that the reconstructed data closely matches the overall distribution of the original data (**Fig.** S2).

#### D Interactive Q&A

##### D.1 Data

Based on the MERFISH Mouse Brain dataset [3], we curate a Q&A dataset covering three tasks: (1) Identifying the top 5 cell types present in a given tissue structure. (2) Listing the top 20 highly expressed genes within a specific tissue structure for a given cell class. (3) Providing a scientific description of the structures within a tissue region.

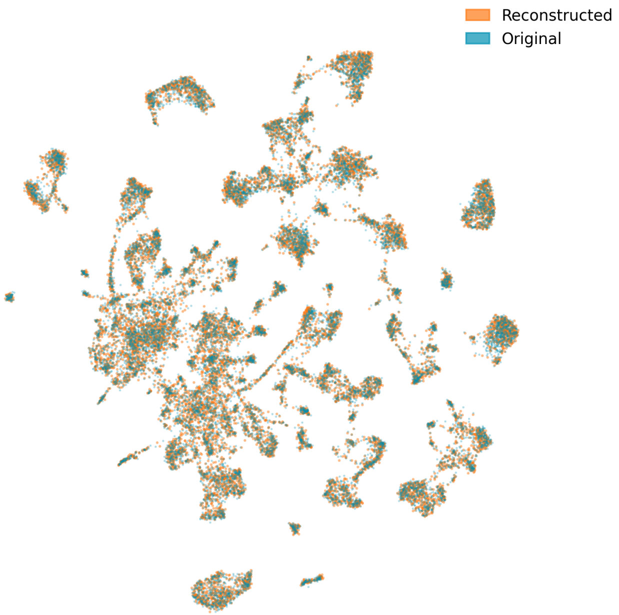


**Figure S2:** UMAP overlay of the original mouse brain data (blue) and the gene expression reconstructed from ranked cell sentences using the top 100 genes (orange). A maximum of 500 cells is sampled per cell type.

For the third task, we generate ground-truth answers by prompting GPT-5 [4] with the tissue region name and all associated structure names within that region. An example for each task is shown in **Fig.** S3.

#### D.2 Results

We split tissue regions into training, validation, and test sets using an $8:1:1$ ratio. Models were fine-tuned separately on TISSUENARRATOR and on the base model for comparison. Fine-tuning on TISSUENARRATOR consistently yields higher performance across tasks, even though TISSUENARRATOR was never trained on the structure names appearing in the Q&A dataset (**Table** S2).

For the second task, the model fine-tuned on TISSUENARRATOR effectively captures region-specific highly expressed genes rather than predicting the same gene set across structures of the same class (**Table** S3).

**Table S2:** Comparison of Overlap Score across three models: Base, fine-tuned on Base, and fine-tuned on TISSUENARRATOR.

| **Task** | **Base** | **Fine-tuned on Base** | **Fine-tuned on TISSUENARRATOR** |
|---|---|---|---|
| Highly expressed genes | 0.0079 | 0.6226 | 0.6554 |
| Top-5 abundant cells | 0.3242 | 0.6564 | 0.6805 |

**Table S3:** Effect of shuffling region-specific information on TISSUENARRATOR performance (Overlap Score).

| **Condition** | **Overlap** |
|---|---|
| Original (with spatial structure) | 0.6485 |
| After shuffling (region information destroyed) | 0.5764 |

##### A. Cell type prediction from spatial region

**Prompt**

```
List the major cell classes present in the Anterior olfactory nucleus region
from the mouse brain (include up to five).
```

**Answer** IT-ET Glut, Vascular, Astro-Epen, OPC-Oligo, Immune

##### B. Highly expressed gene prediction from spatial region

**Prompt**

```
List the top 20 highly expressed genes in the Astro-Epen cell class of the
Accessory olfactory bulb region from the mouse brain. Provide your answer as
a ranked list of gene names separated by commas.
```

**Answer** Gja1, Mfge8, Acsbg1, Gpr37l1, S1pr1, Ntsr2, Aqp4, Pla2g7, Serpine2, Pou3f3, Fam107a, Gfap, Grin2c, Slc7a10, Aldh1l1, Gprc5b, Id4, Itih3, Sox9, Sox2

##### C. Spatial region summarization

**Prompt**

```
Provide a concise scientific summary of the brain division 'cranial nerves'
from the mouse brain. Describe its overall functional role and highlight
representative structures and their significance.
```

**Answer** The cranial nerves are a set of peripheral pathways that connect the brain and brainstem to the sensory organs, muscles, and autonomic targets of the head and neck, with the vagus extending this control to thoracic and abdominal viscera; collectively they carry general and special sensory inputs, somatic and branchial motor outputs, and parasympathetic (visceromotor) signals, enabling perception (vision, olfaction, hearing, balance), precise motor control (eye, facial, oral–pharyngeal), homeostatic reflexes, and autonomic regulation. Among representative examples, the optic nerve (II) conveys high-fidelity visual information from retinal ganglion cells to central visual pathways and, though named a nerve, is a CNS tract myelinated by oligodendrocytes, underscoring the cranial system's tight integration with the brain. The trigeminal nerve (V) ...

**Figure S3:** Examples of spatial question–answering tasks, illustrating **A.** cell type prediction, **B.** highly expressed gene prediction, and **C.** spatial region summarization.

### E Baseline Implementation Details

For a fair comparison with the conditioned version of TISSUENARRATOR, we introduce the following modifications to each baseline method. **Nearest Neighbor**: modified to select the nearest neighbor belonging to the same class as the target cell in the prompt. **Class Mean**: no change is required, as it already incorporates class information. **Nicheformer**: the neighbor embedding is concatenated with an additional embedding representing the class of the target cell, and the combined representation is passed to an MLP prediction head. **C2S-Scale** and **Base**: the prompt is augmented with metadata describing the target cell.

### F Ablation Study

We conduct an ablation study to evaluate the contribution of different components in TISSUENARRATOR. All experiments are performed under the cross-type generation setting using the conditioned generation mode. The following model variants are compared:

1. **No meta**: metadata describing the target cell is removed.
2. **No pos**: spatial coordinates $(X, Y)$ are removed.
3. **Random traversal**: spatial sentences are constructed using a random traversal rule during training.
4. **No neighbor**: neighbor information is excluded during inference.

As shown in **Table** S4, each component contributes to the overall performance of TISSUENARRATOR. Among them, neighbor information and metadata are the most influential.

**Table S4:** Ablation results showing Overlap@50 and NDCG@50 for TISSUENARRATOR and its variants.

| Model Variant | Overlap@50 | NDCG@50 |
|---|---|---|
| TISSUENARRATOR (full model) | 0.420 | 0.853 |
| No meta | 0.229 | 0.644 |
| No pos | 0.407 | 0.841 |
| Random traversal | 0.417 | 0.852 |
| No neighbor | 0.337 | 0.764 |
